# Fast oscillations >40Hz localize the epileptogenic zone: an electrical source imaging study using high-density electroencephalography

**DOI:** 10.1101/2020.03.02.973602

**Authors:** Tamir Avigdor, Chifaou Abdallah, Nicolás von Ellenrieder, Tanguy Hedrich, Annalisa Rubino, Giorgio Lo Russo, Boris Bernhardt, Lino Nobili, Christophe Grova, Birgit Frauscher

## Abstract

**Objective:** Fast Oscillations (FO) >40 Hz are a promising biomarker of the epileptogenic zone (EZ). Evidence using scalp electroencephalography (EEG) remains scarce. We assessed if electrical source imaging of FO using 256-channel high-density EEG (HD-EEG) is useful for EZ identification.

**Methods:** We analyzed HD-EEG recordings of 10 focal drug-resistant epilepsy patients with seizure-free postsurgical outcome. We marked FO candidate events at the time of epileptic spikes and verified them by screening for an isolated peak in the time-frequency plot. We performed electrical source imaging of spikes and FO within the Maximum Entropy of the Mean framework. Source localization maps were validated against the surgical cavity.

**Results:** We identified FO in five out of 10 patients who had a superficial or intermediate deep generator. The maximum of the FO maps was localized inside the cavity in all patients (100%). Analysis with a reduced electrode coverage using the 10-10 and 10-20 system showed a decreased localization accuracy of 60% and 40% respectively.

**Conclusions:** FO recorded with HD-EEG localize the EZ. HD-EEG is better suited to detect and localize FO than conventional EEG approaches.

**Significance:** This study acts as proof-of-concept that FO localization using 256-channel HD-EEG is a viable marker of the EZ.

**Highlights:**

- Fast oscillations > 40Hz are able to correctly localize the epileptogenic zone.
- HD-EEG is superior in detection and localization of fast oscillations compared to conventional EEG approaches.
- Presence of fast oscillations on the scalp might point to a superficial epileptic generator.

## 1. INTRODUCTION

Epilepsy is a chronic condition characterized by recurrent seizures accompanied by negative impact on quality of life (Hinnell et al., 2010). A significant number of 30% of patients with focal epilepsy are drug-resistant, and these numbers did not change over the past 30 years despite the development of multiple new antiepileptic drugs (Chen et al., 2018). The therapy of choice for focal drug-resistant epilepsy is epilepsy surgery (Vakharia et al., 2018). The aim of epilepsy surgery is to remove the epileptogenic zone (EZ) defined as the area of cortex that needs to be removed to achieve seizure freedom (Rosenow and Lüders, 2001). In current practice, the seizure-onset zone (SOZ) is used as the main proxy marker for the EZ. A sustained seizure-free condition is currently, however, obtained in only 50% of carefully selected patients (Krucoff et al., 2017; Mohan et al., 2018; West et al., 2019). This is likely due to inaccurate localization of the EZ or a network involvement larger than initially expected (Englot, 2018). This underlines the need to develop new markers and localization techniques for better identification of the EZ and hence improved surgical outcomes.

Recently, Fast Oscillations (FO) have been identified as promising novel interictal marker for the EZ (Frauscher et al., 2017). Most evidence comes from intracranial electroencephalography (iEEG) suggesting that resection of areas with high FO rates is associated with good surgical outcome, and that presence of FO after resection is predictive of postsurgical seizure relapse (Frauscher et al., 2017; Höller et al., 2015; van ‘t Klooster et al., 2017; Jacobs et al., 2018; Tamilia et al., 2018). In contrast, evidence from non-invasive methods is rather scarce (Thomschewski et al., 2019), and combination of FO detection with source localization was mainly obtained from magnetoencephalography (MEG). MEG studies from different centers suggest that it is possible to detect and then localize isolated FO events > 40 Hz from interictal MEG recordings with satisfactory data quality (von Ellenrieder et al., 2016; van Klink et al., 2016a; Papadelis et al., 2016). Four studies validated FO localization against the postsurgical cavity in good outcome patients as gold standard for the EZ; they pointed towards usefulness of FO for delineating the EZ (van Klink et al., 2017; Velmurugan et al., 2019; Yin et al., 2019; Tamilia et al., 2020).

In contrast to MEG, high-density EEG (HD-EEG) would have the great advantage that it is easy to maintain and more affordable given the high cost of operating a MEG device. From a technical point of view, however FO recording and localization using HD-EEG has even more challenges to overcome. The main hurdle is a more complex solution of the forward problem needed for source localization given the necessity of assessing accurately the conductivity of the head tissues, especially the skull (Goldenholz et al., 2009; de Munck et al., 2012; Hari and Puce, 2017; Ilmoniemi and Sarvas, 2019). There is first evidence that FO can be detected also in the scalp EEG (Andrade-Valenca et al., 2011; Kobayashi et al., 2011; von Ellenrieder et al., 2012; Zelmann et al., 2014; van Klink et al., 2016b; Pizzo et al., 2016). Some of these studies showed concordance with the area of the seizure-onset zone (SOZ) as determined by intracranial EEG or the resection cavity in patients with good seizure outcome (Kuhnke et al., 2018; Kuhnke et al., 2019; Tamilia et al., 2020). Moreover, it was shown that presence of scalp FO is associated with the depth of the epileptic generator, with FO being present in case of a generator located in more superficial cortical regions (Cuello-Oderiz et al., 2017). Finally, it was suggested that scalp EEG might be superior for the detection of FO due to higher rates of FO detected in scalp EEG compared to MEG (van Klink et al., 2019, Tamilia et al., 2020). Given the small size of the generators of FO (von Ellenrieder et al., 2014), HD-EEG might then be superior to standard EEG. So far, there is only one study reporting scalp FO using HD-EEG with 128 electrodes in the sensor space (Kuhnke et al., 2018).

In this proof-of concept study, we assessed the feasibility of detection and localization of FO in epileptic patients using a HD-EEG array of 256 electrodes. We examined whether localization of FO >40 Hz using HD-EEG is capable of delineating the EZ (Rosenow and Lüders, 2001). We then compared results to standard spike source localization and to conventional EEG approaches using the 10-10 or 10-20 EEG system. For validation, we opted to use the surgical cavity in postsurgically seizure-free patients (Engel Ia) with a follow-up duration of > 2 years as best approximation of the EZ. We opted to not use the intracranially identified SOZ as alternative validation standard, given that a considerable number of patients do not become seizure-free after removal of the SOZ (Krucoff et al., 2017).

## 2. METHODS

### 2.1 Subject selection

A total of ten patients with seizure-free outcome > 2 years (Engel 1A (Engel, 1993)) after epilepsy surgery procedure were selected from the HD-EEG database of the Claudio Munari Epilepsy Center in Milan between 2015-2016. Selection criteria were presence of >10 epileptic spikes during HD-EEG recording, presence of non-rapid eye movement (NREM) sleep in order to have the least artifacts possible for FO detection (Zijlmans et al., 2017), absence of previous surgery, availability of electrode positions and magnetic resonance imaging (MRI) coregistration and satisfactory data quality. Figure 1 presents the flowchart for patient selection. Note that data quality was a negligible reason for exclusion. Patient demographic and clinical information is provided in Table 1. The study conforms to the Declaration of Helsinki and was approved by Niguarda Hospital in Milan, Italy. A written informed consent was signed by all patients prior to study participation.

**Figure 1.**
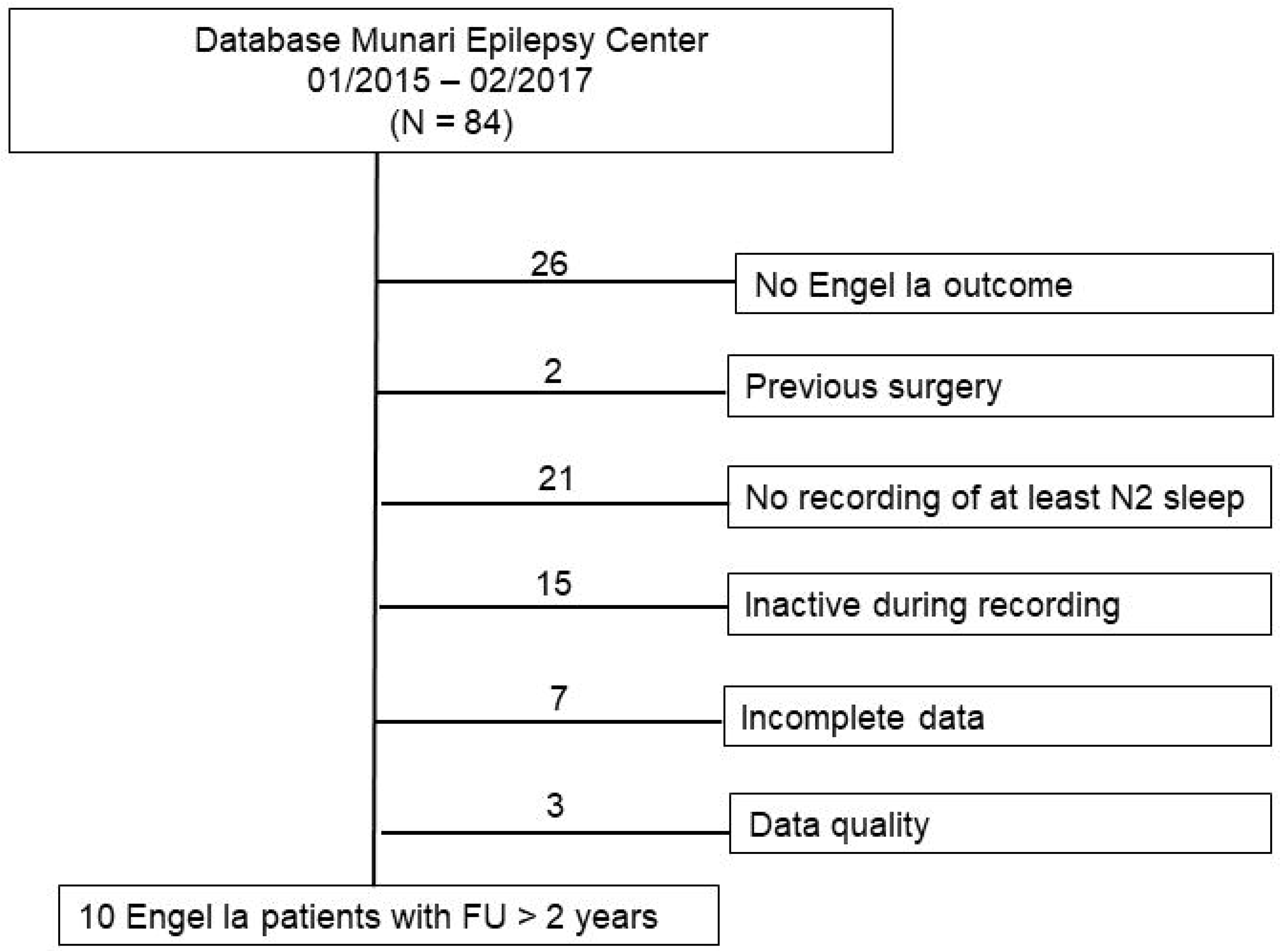
Flowchart describing the selection process from the complete high-density electroencephalogram database of surgical epilepsy patients comprising 87 patients to the final 10 patients analyzed in this study.

**Table 1.**
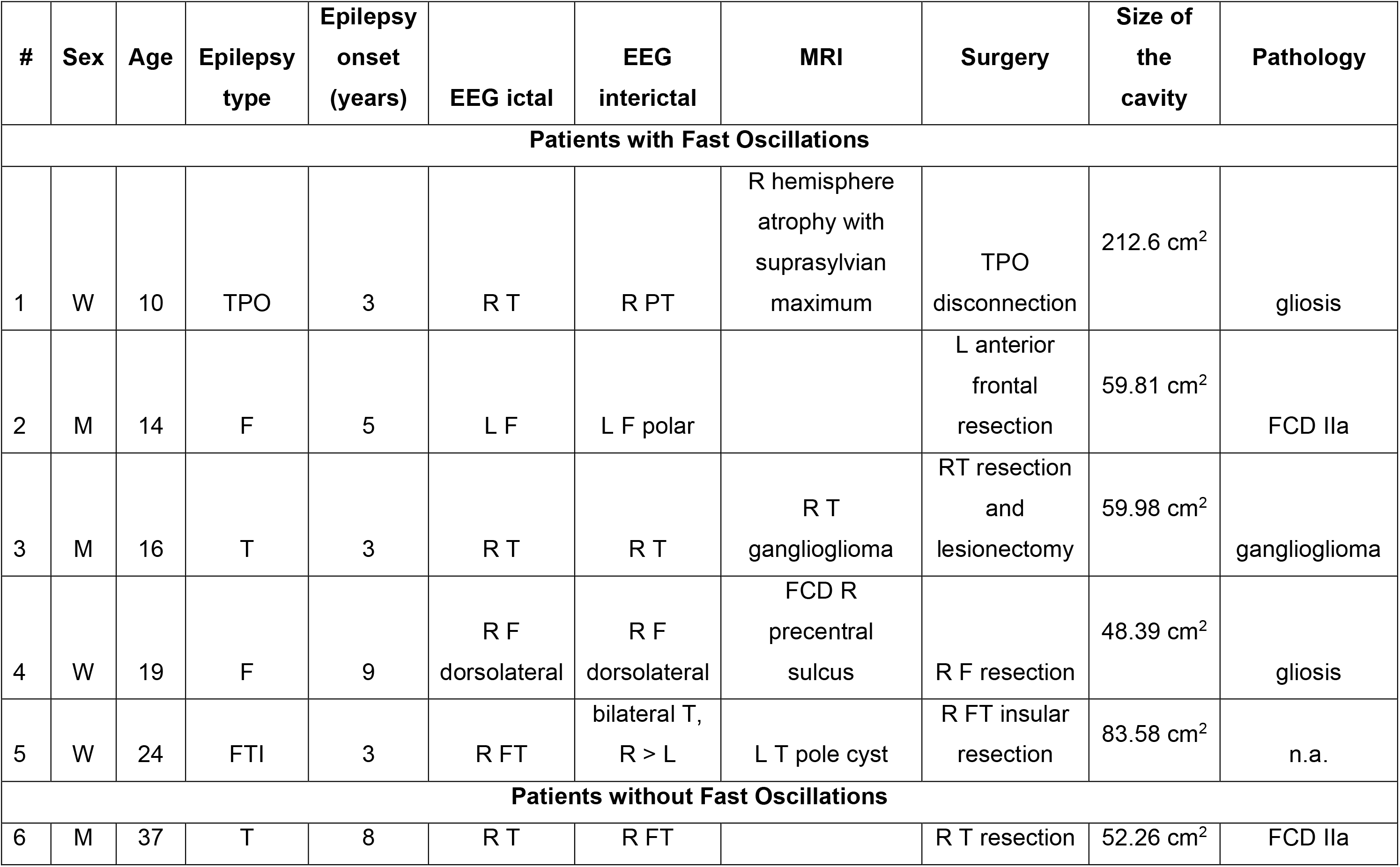
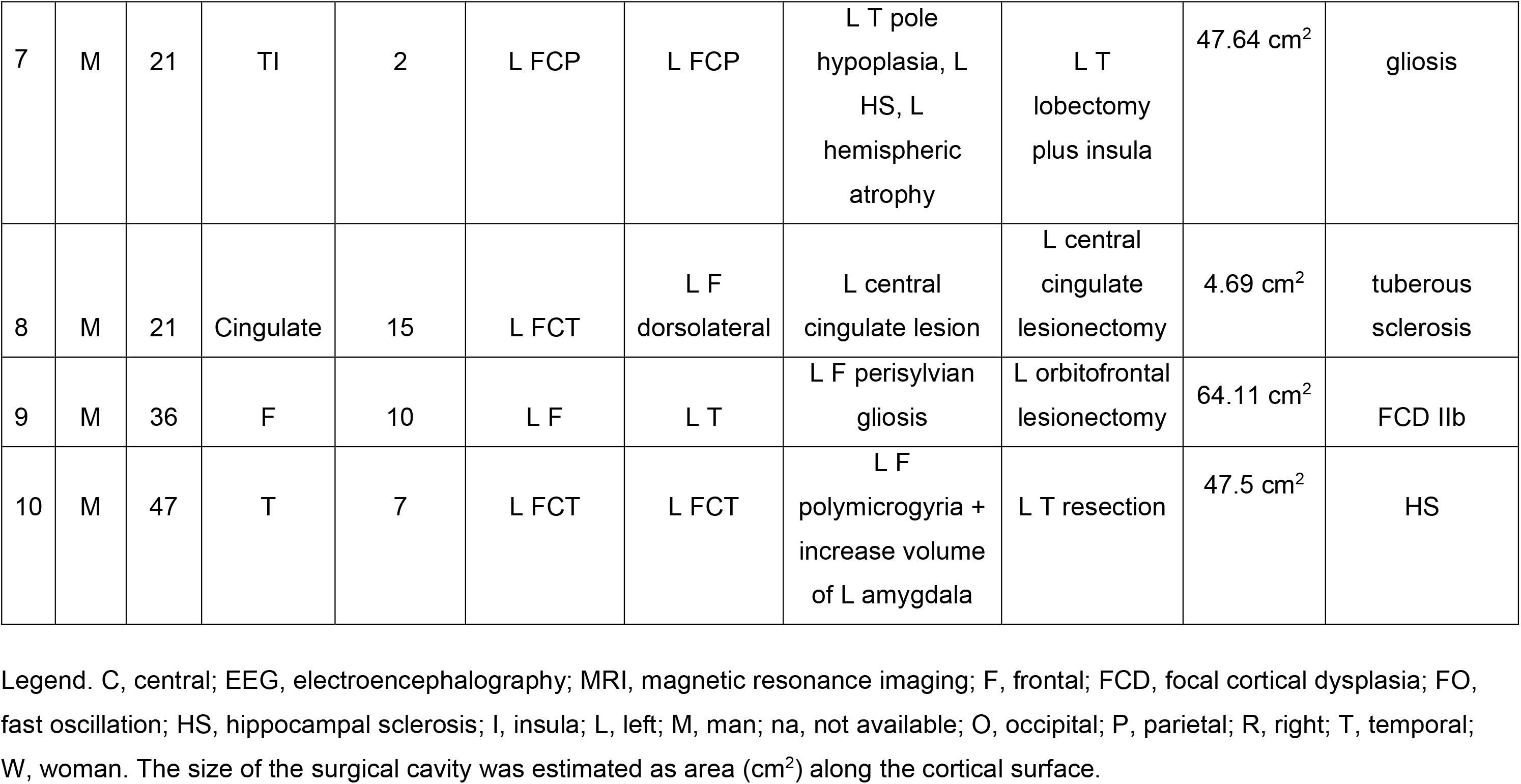
Demographic and clinical information of the study sample.

| # | Sex | Age | Epilepsy type | Epilepsy onset (years) | EEG ictal | EEG interictal | MRI | Surgery | Size of the cavity | Pathology |
| --- | --- | --- | --- | --- | --- | --- | --- | --- | --- | --- |
| **Patients with Fast Oscillations** |  |  |  |  |  |  |  |  |  |  |
| 1 | W | 10 | TPO | 3 | R T | R PT | R hemisphere atrophy with suprasylvian maximum | TPO disconnection | 212.6 cm<sup>2</sup> | gliosis |
| 2 | M | 14 | F | 5 | L F | L F polar |  | L anterior frontal resection | 59.81 cm<sup>2</sup> | FCD IIa |
| 3 | M | 16 | T | 3 | R T | R T | R T ganglioglioma | RT resection and lesionectomy | 59.98 cm<sup>2</sup> | ganglioglioma |
| 4 | W | 19 | F | 9 | R F dorsolateral | R F dorsolateral | FCD R precentral sulcus | R F resection | 48.39 cm<sup>2</sup> | gliosis |
| 5 | W | 24 | FTI | 3 | R FT | bilateral T, R > L | L T pole cyst | R FT insular resection | 83.58 cm<sup>2</sup> | n.a. |
| **Patients without Fast Oscillations** |  |  |  |  |  |  |  |  |  |  |
| 6 | M | 37 | T | 8 | R T | R FT |  | R T resection | 52.26 cm<sup>2</sup> | FCD IIa |
| 7 | M | 21 | TI | 2 | L FCP | L FCP | L T pole<br>hypoplasia, L<br>HS, L<br>hemispheric<br>atrophy | L T<br>lobectomy<br>plus insula | 47.64 cm<sup>2</sup> | gliosis |
| 8 | M | 21 | Cingulate | 15 | L FCT | L F<br>dorsolateral | L central<br>cingulate lesion | L central<br>cingulate<br>lesionectomy | 4.69 cm<sup>2</sup> | tuberous<br>sclerosis |
| 9 | M | 36 | F | 10 | L F | L T | L F perisylvian<br>gliosis | L orbitofrontal<br>lesionectomy | 64.11 cm<sup>2</sup> | FCD IIb |
| 10 | M | 47 | T | 7 | L FCT | L FCT | L F<br>polymicrogyria +<br>increase volume<br>of L amygdala | L T resection | 47.5 cm<sup>2</sup> | HS |

Legend. C, central; EEG, electroencephalography; MRI, magnetic resonance imaging; F, frontal; FCD, focal cortical dysplasia; FO, fast oscillation; HS, hippocampal sclerosis; I, insula; L, left; M, man; na, not available; O, occipital; P, parietal; R, right; T, temporal; W, woman. The size of the surgical cavity was estimated as area (cm<sup>2</sup>) along the cortical surface.

### 2.2 HD-EEG data acquisition and preprocessing

HD-EEG was recorded using a 256-electrode EEG system (Electrical Geodesic Inc., EGI, now Magstim EGI, Eden Prairie, MN, USA) with a sampling rate of 500 Hz and hardware filter settings of 0.3 Hz for the high pass and 200 Hz for the low pass filter. The recordings were performed with long-term EEG nets using gel and lasted approximately 1.5 h; this duration was chosen to enable the patient to fall asleep. The impedance of the selected electrodes was kept under 1 kΩ. Cz was the recording reference for this study. For analysis, we created an average referential montage.

HD-EEG processing was performed with the Brainstorm software package (Tadel et al., 2011). For interictal spike detection and analysis, preprocessing included 0.3–70 Hz band-pass filtering and direct current correction (baseline window from −1000 ms to−500 ms before the marked spikes). No filter was applied for FO events.

The EEG sensor positions were estimated using digitalization with a SofTaxicOptic system (EMS s.r.l., Bologna, Italy). A linear coregistration with a pre-implant MRI (Achieva 1.5 T, Philips Healthcare) was performed. The digitized positions of the electrodes were then coregistered to the scalp surface segmented from the anatomical MRI of each patient, using a surface matching algorithm within the Brainstorm software. The accuracy of the resulting registration was then verified visually. The HD-EEG electrodes located on the face and on the neck (~40 channels) were excluded for further analysis in order to avoid artifacts caused by muscle or poor-contact electrodes (Hedrich et al., 2017).

### 2.3 Interictal event marking

Spikes were marked at their peak by an epileptologist (CA). All spike events occurring during N1, N2 or N3 sleep were screened for FO candidate events by a second epileptologist (BF). FO were defined as at least four oscillations clearly standing out of the background EEG in the gamma (40–80 Hz) and ripple (80-160 Hz) frequency band in the same electrodes as the spike that was marked at that time. FO candidate events were verified using a time-frequency representation screened for an isolated peak within the FO band (gamma or ripple) with no other visible peaks in the time frequency (TF) plot in the same threshold within 0.5 s before and 0.5 s after (Figure 2). TF representations were computed using the Morlet wavelet with a central frequency of 1 Hz and a window duration of 3s and plotted as a power scale. We used this two-step approach in order to avoid misclassification of the filtering effect as “true” FO. Given the low signal to noise ratio (SNR) of FO, we attempted source localization only in patients with a minimum number of 5 FO in either the 40-80 or 80-160 Hz window.

**Figure 2.**
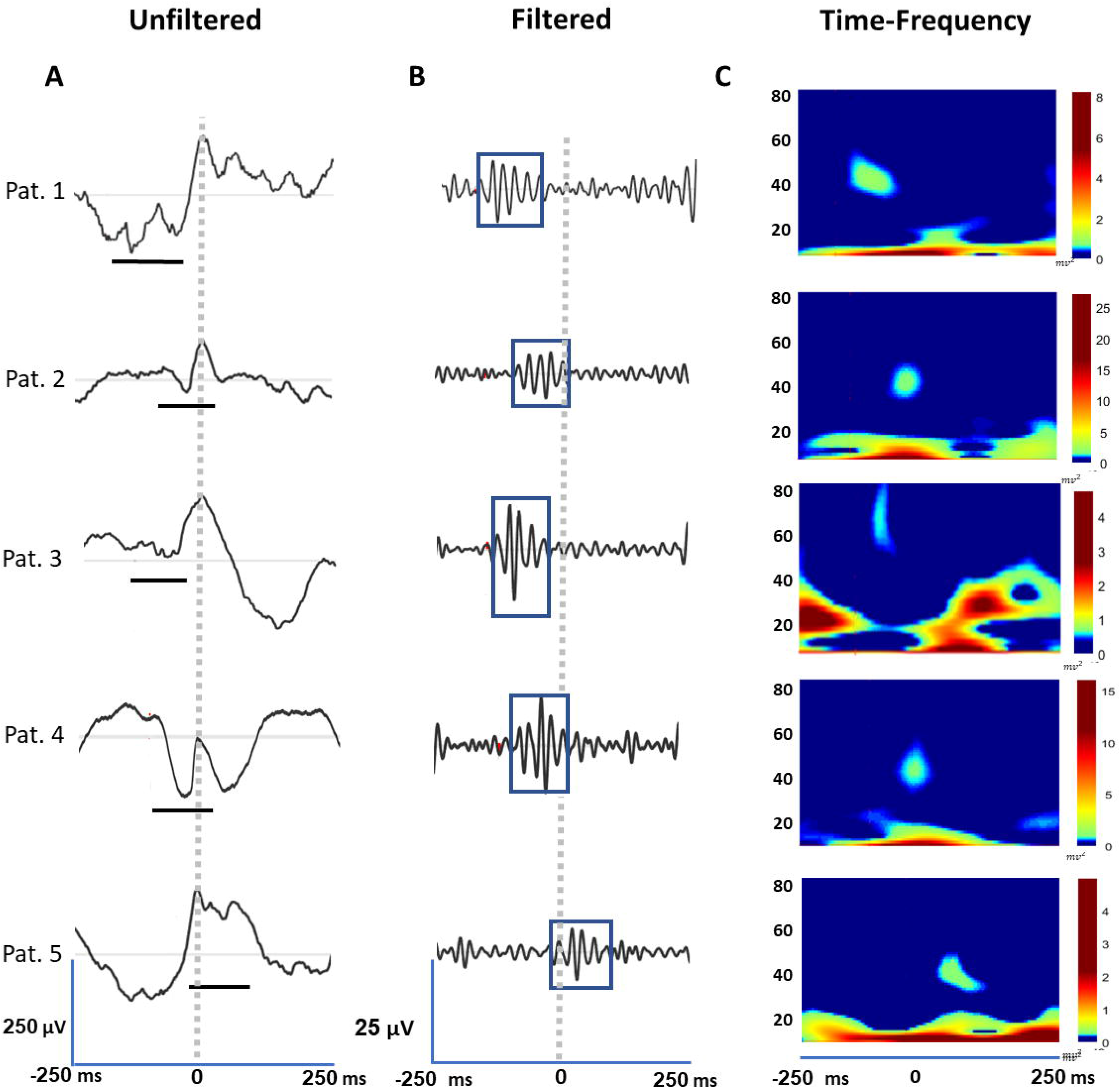
Detection of fast oscillations (FO) using scalp high-density electroencephalogram (HD-EEG). This figure provides examples of FO for each patient. (A) unfiltered signal showing the epileptic spike with the FO underlined in black (B) filtered single (40-80 Hz) demonstrating the FO event over the background (C) time frequency plot showing the isolated peak in the 40-80 Hz frequency band atop of the spike. Time scale is set to the peak of the event and 125 ms before and after are given.

### 2.4 Electrical source imaging

The 1.5T anatomical MRI was segmented, and the cortical surface was obtained using the FreeSurfer software package (v=6.0, http://surfer.nmr.mgh.harvard.edu). The EEG forward problem was solved using the boundary element method (BEM) (Kybic et al., 2006). The gain matrix was calculated using a 3-layer BEM model for brain, skull and scalp (conductivity of 0.33, 0.0165, 0.33 S/m respectively) using OpenMEEG (Gramfort et al., 2010) implemented in the Brainstorm software. The inverse problem was solved using the Maximum Entropy on the Mean (MEM) method (Amblard et al., 2004). MEM is a nonlinear distributed inverse problem method, for which the prior model is built using a data-driven parcellation (DDP) technique in order to cluster the cortical surface into K parcels (Lapalme et al., 2006). To do so, we used the multivariate source pre-localization (MSP) method (Mattout et al., 2005) which is a projection method that estimates a coefficient, which characterizes the possible contribution of each dipolar source to the data. We make use of it in the MEM reference model (or prior model), in which a hidden variable is associated to each parcel in order to model the probability of the parcel to be active or to be switched off. This method will be referred to as the coherent MEM (cMEM), which was carefully evaluated for its ability to recover the spatial extent of the underlying generators (Chowdhury et al., 2013; Chowdhury et al., 2016; Hedrich et al., 2017; Pellegrino et al., 2018, Pellegrino et al., 2020). The wavelet based MEM extension (wMEM) of the MEM framework (Lina et al., 2014) decomposes the signal in a discrete wavelet basis before performing MEM source localization on each TF box. Therefore, wMEM is particularly suited to localize oscillatory patterns, as evaluated with realistic simulations, but also when localizing FO (von Ellenrieder et al., 2016), oscillatory patterns at seizure onset (Pellegrino et al., 2016) or resting state ongoing oscillations (Aydin et al., 2020). These two different variations of MEM were used in this study: wavelet MEM (wMEM) was adopted to localize FO, whereas cMEM (cMEM) was adopted to localize spikes. We used the standard settings of the wMEM and cMEM as provided in Brainstorm (Chowdhury et al., 2013; Lina et al., 2014; https://github.com/multifunkim/best-brainstorm), except that for wMEM we set the amount of vanishing moments for the Daubechies wavelet to 8. The use of 8 vanishing moments instead of the default 4 was done in order to capture the complexity of FO. The baseline to model the diagonal noise covariance matrix for both wMEM and cMEM was chosen for each patient visually during an artifact- and spike-free 2-second period. wMEM was performed on the marked duration of the FO by selecting the only TF box exhibiting the largest amount of energy as recommended (von Ellenrieder et al., 2016). To select which TF boxes were considered as localizable using the wMEM framework, a global-spatial wavelet power was first computed for each time-frequency box, by summing up the energy of each wavelet coefficient over all the sensors. Total power was then estimated by considering all frequencies (scales) and time samples of a specific segment of the duration of the FO. TF boxes that were not contributing to 99% of the total cumulative power, within the selected window, were considered as too low SNR and therefore non localizable. FOs, for which no localizable TF box was identified within the frequency band of interest, were labelled as non localizable FO events. The spike map was computed for −50 to +50 ms from the spike peak using cMEM. Spikes were filtered using a 0.3-70Hz finite impulse response filter prior to cMEM localization; for FO no filter was applied as wMEM acts a filter which is chosen by a specific frequency band, i.e. 40-80 Hz or 80-160 Hz.

### 2.5 Epileptic spike and FO consensus maps

In order to take into account reliability and reproducibility of FO or spikes, we applied the concept of consensus maps of source localization (Chowdhury et al., 2018). In order to estimate these consensus maps for every patient, we first applied cMEM or wMEM to generate source maps of every single event, FO or spikes. In a second step, we estimated a similarity index between all single event source maps (FO or spikes), based on spatio-temporal correlation around the peak of the event The source maps were then clustered using a hierarchical clustering approach (Ward’s hierarchical clustering) followed by thresholding of the dendrogram to obtain the optimal number of clusters. The cluster exhibiting most events was selected and the consensus map was finally obtained by averaging all the single event maps of this cluster. This consensus approach was considered instead of simple averaging of the event followed by one source localization, in order to enhance reliability between source localization results and reducing the influence of more noisy maps. We previously carefully demonstrated the robustness of this approach when considering either EEG source imaging, MEG source imaging or EEG-MEG fusion source imaging (Chowdhury et al., 2018).

### 2.6 Comparing HD-EEG against conventional EEG approaches

Our main results used the 256-electrode montage (after removal of ~40 face and neck artifactual electrodes per patient). We then compared our results to the standard 10-10 system (73 electrodes) and the 10-20 system (25 electrodes). The 10-10 and 10-20 system was approximated from the full montage as per the recommendations of EGI geodesic systems (see Figure 3 for selected electrodes). All FO were localized using these three montages and results were compared using the same metrics (see below).

**Figure 3.**
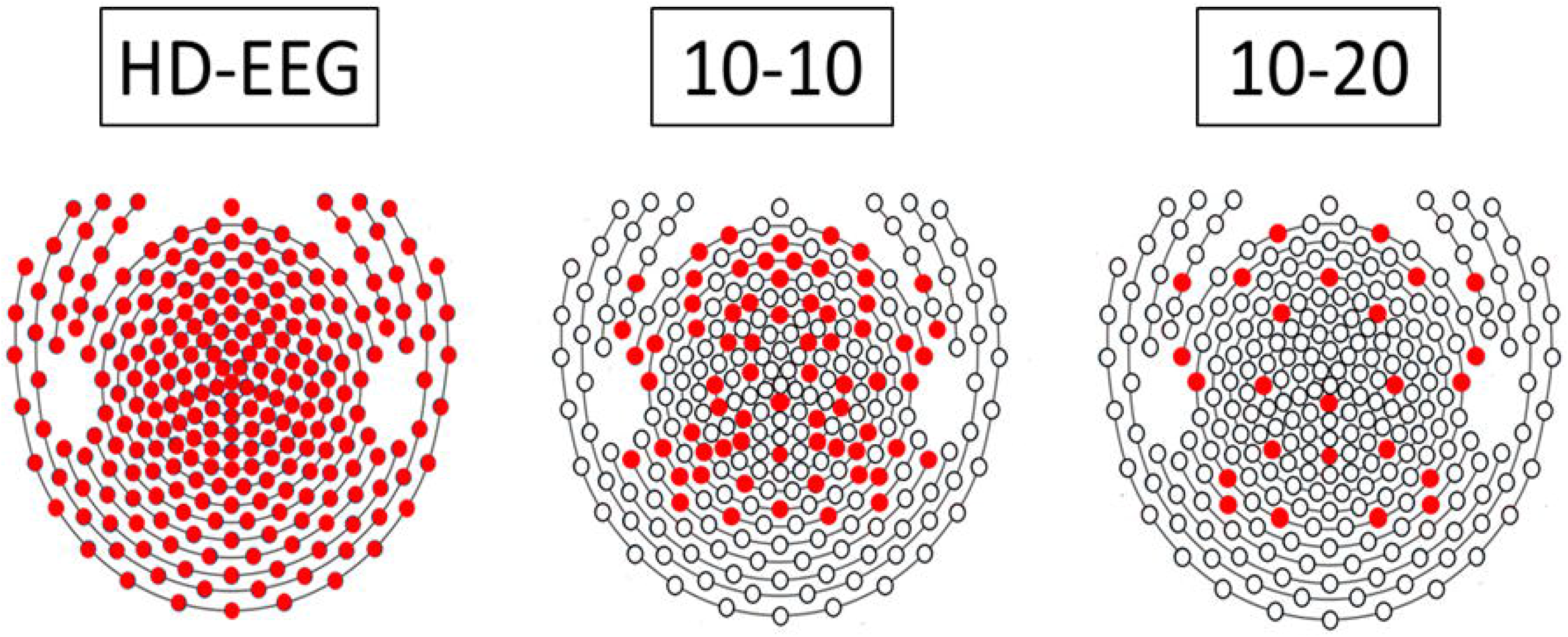
Illustration of the distribution of electrodes in the 256 channel high-density electroencephalography (HD-EEG) system versus the standard 10-10 system (73 electrodes) and the 10-20 system (25 electrodes). Note that the 10-10 and 10-20 systems were approximated from the full HD-EEG montage as per the recommendations of the EGI geodesic systems manual.

### 2.7 Generator depth

The depth of the generator was assessed on the basis of all available clinical information. The generator depth was divided into 3 categories: superficial, intermediate and deep as done in previous work (Cuello-Oderiz et al., 2017). Superficial was considered as involving the neocortex adjacent to the skull including the bottom of the sulcus; deep was considered as involving the medial aspects of the frontal, parietal, occipital, and temporal lobes and temporo-occipital basal regions; and intermediate as generators found in regions not fitting the above two categories.

### 2.8 Evaluation Metrics & Statistics

All final spikes and FO consensus maps were tresholded at 50% of the maximum reconstructed intensity for both visualization and statistical purposes. We previously demonstrated that MEM source localization results provide maps with high contrast, therefore results on the spatial extent are relatively stable within a large range of detection threshold, as opposed to other standard source localization techniques (Chowdhury et al., 2018; Pellegrino et al., 2018; Pellegrino et al., 2020). In order to assess the success or failure of the localization, we considered the surgical cavity as a reference. Since all selected patients are > 2 year postsurgically seizure-free (Engel 1a), we are sure that the presumed EZ was indeed localized within the cavity. The surgical cavity was fitted as a surface-based region of interest (scout) on the presurgical cortical surface considered as our brain source model (Supplementary Figure 1). This was done visually using the post-surgical MRI co-registered to the intact cortical surface of the pre-surgical MRI, using the Brainstorm software. The coregistration between presurgical and postsurgical MRI was obtained with a 6 parameter rigid body coregistration using the MINC toolkit (https://bic-mni.github.io/). All evaluation metrics were then estimated using this scout of the surgical cavity as our clinical reference. We assessed the following validation metrics: the minimum Distance Localization Error (Dmin), Spatial Dispersion (SD), the Spatial Map Intersection (SMI), and the SNR. For each of the validation metrics described above, we considered only the consensus localization map as defined as the average localization obtained from the cluster with exhibiting the largest number of single events for both spikes and FO.

- Dmin: the minimum distance localization error was computed as the Euclidean distance in mm from the maximum of the map to the closest vertex belonging to the cavity. Whenever this maximum was located inside the cavity, Dmin was set to 0 mm.
- Spatial Dispersion (SD): the SD metric measured the spatial spread (in mm) of the localization around the Ground Truth considered here as the surgical cavity. It was computed as the root mean square of the distance from the cavity weighted by the energy of the source localization map on each vertex.

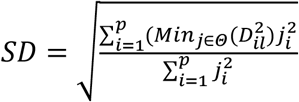 Where Θ denoted the surgical cavity, and *j_i_*, is the amplitude results of the MEM solver (Cmem or wMEM) for each dipolar source *i*, while 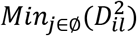 function provides the minimum Euclidean distance between the dipolar source i to the closest vertex within the surgical cavity Θ.
- Spatial Map Intersection (SMI): SMI was defined as the percentage of the thresholded localization map that was falling within the surgical cavity region of interest, such that if all the thresholded localization was inside the scout then it equals 100% intersection.

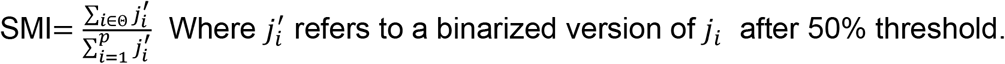
- Signal to noise ratio (SNR): the SNR was calculated using the mean amplitude at the time of the event divided by the standard deviation of a baseline period of 10 ms long for 200ms prior to the event, the calculation was done on the filtered signal for each type of event (0.3-70Hz for spikes and 40-80Hz of 80-160 Hz for FO) in accordance to the localization filters. For spikes the SNR calculation was made on the average spike. For FO, SNR was calculated for each event and then averaged over all events of each patient, followed by averaging over all patients to reach the final SNR. We used the Welch’s t-test to compare the different metrics. The Fisher exact test was used to assess the distribution of generator depth categories between patients with and without FO.

## 3. RESULTS

### 3.1 Detection of FO

FO in the gamma band (40-80 Hz) were found in five out of ten patients (Table 2). FO in the ripple band (>80Hz) did not reach the minimum of 5 requested events per patient. The mean number of FO > 40 Hz was 12.2±3.9 (range, 10-20) with a mean SNR of 3.4±1.4. Figure 2 shows representative examples of FO of the five patients in the unfiltered signal, the filtered signal, as well as the corresponding time frequency plot. FO preceded spikes in 78.8% of cases with an onset of 0.1± 0.03 s before the peak of the spike. Spikes were present in all ten patients. The median number of spikes was 41 (range, 12-303) with a mean SNR of 9.3±2.7. As expected, the SNR was significantly lower in FO than in spikes (p<0.05).

**Table 2.** Depth of the epileptic generator, epileptic spikes, and fast oscillations (FO).

| # | Depth of generator | # Spikes localized | # FO localized | # of Spikes in the consensus map | # of FO in the consensus map | Localizable FO main frequency (Hz) range |
| --- | --- | --- | --- | --- | --- | --- |
| **Patients with Fast Oscillations** |  |  |  |  |  |  |
| **1** | superficial | 303 | 10 | 164 | 5 | 42-65 |
| **2** | superficial | 258 | 11 | 101 | 10 | 41-73 |
| **3** | intermediate | 155 | 11 | 69 | 6 | 65-80 |
| **4** | superficial | 12 | 20 | 8 | 6 | 40-55 |
| **5** | superficial | 20 | 9 | 10 | 6 | 42-80 |
| **Patients without Fast Oscillations** |  |  |  |  |  |  |
| **6** | deep | 92 |  | 33 |  |  |
| **7** | deep | 42 |  | 23 |  |  |
| **8** | deep | 40 |  | 20 |  |  |
| **9** | intermediate | 34 |  | 22 |  |  |
| **10** | deep | 40 |  | 16 |  |  |

For each patient the number of localizable epileptic spikes and fast oscillations (FO) are presented. The depth of the generator is given. It is based on the available clinical information and rated by an epileptologist. Note that FOs > 80Hz were found in 8/10 patients. As their number did not reach five or more FOs > 80 Hz, they were not included in the analysis.

### 3.2 Electrical source imaging

FO sources were localized in 5 out of 5 patients inside the surgical cavity (Figure 3) resulting in a minimum distance Dmin of 0 mm. FO source maps were localized within the surgical cavity with an SD of 9.4±3.2 mm and a SMI of 62.0±15.0%. Spike sources localized inside the surgical cavity in 9 out of 10 patients (Figure 4, Supplementary Figure 2) with a Dmin of 6.5±20.4 mm. Source maps were localized within the surgical cavity with an SD of 6.2±10.9 mm and a SMI of 76.0±30.0%. The spread outside the cavity as well as the activation intersection with the surgical cavity did not differ significantly between FO and spikes (SD: p=0.17; SPI: p=0.09). For further details see Figure. 5.

**Figure 4.**
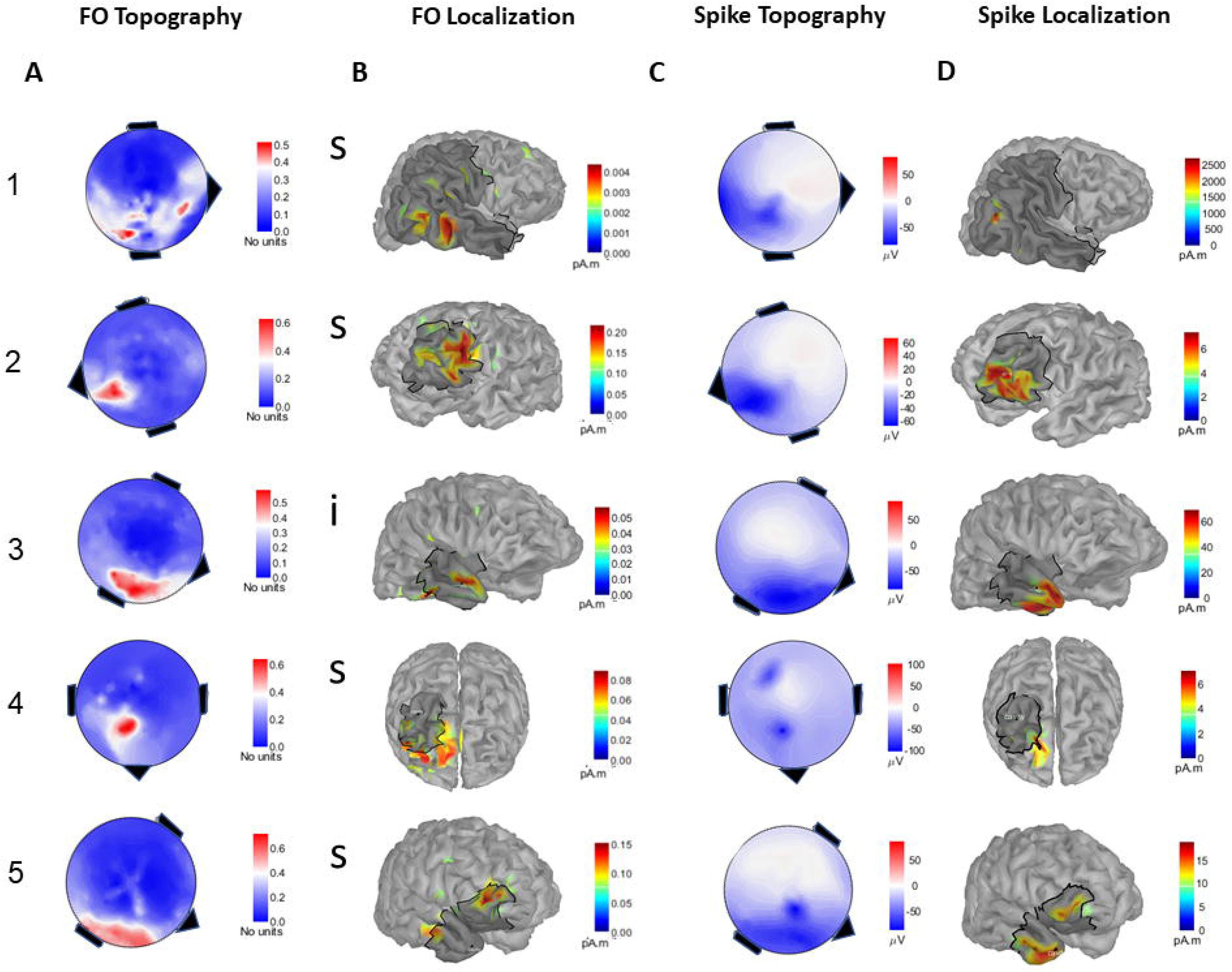
Results of electrical source imaging for fast oscillations (FO) and spikes. FO and spikes were source localized using the Maximum Entropy on the Mean (MEM) method (wavelet MEM for FO and coherent MEM for spikes) in patients who showed > 5 FO. The surgical cavity was fitted on the brain model and was marked as grey area. (A) mean scalp topography of the consensus map (defined as cluster exhibiting most single events) of FO for each patient. Power topography maps were averaged together for the peak of the event time course over the peak frequency as visualized by the time frequency plot (B) consensus map of electrical source imaging (wMEM) of selected FO for each patient (C) mean scalp topography of the consensus map of spikes for each patient (D) consensus map of electrical source imaging (cMEM) of selected spikes for each patient. Please note that scalp topographies (column A and C) are presented following the same orientation as the corresponding source maps. All source localization results are presented using a color map scaled to the maximum reconstructed intensity of the corresponding map and thresholded at 50% of their maximum value. The current amplitude of sources of FO was as expected several orders of magnitude lower than that of spike sources. Patient 1’s large cavity is due to a disconnecting surgery.

**Figure 5.**
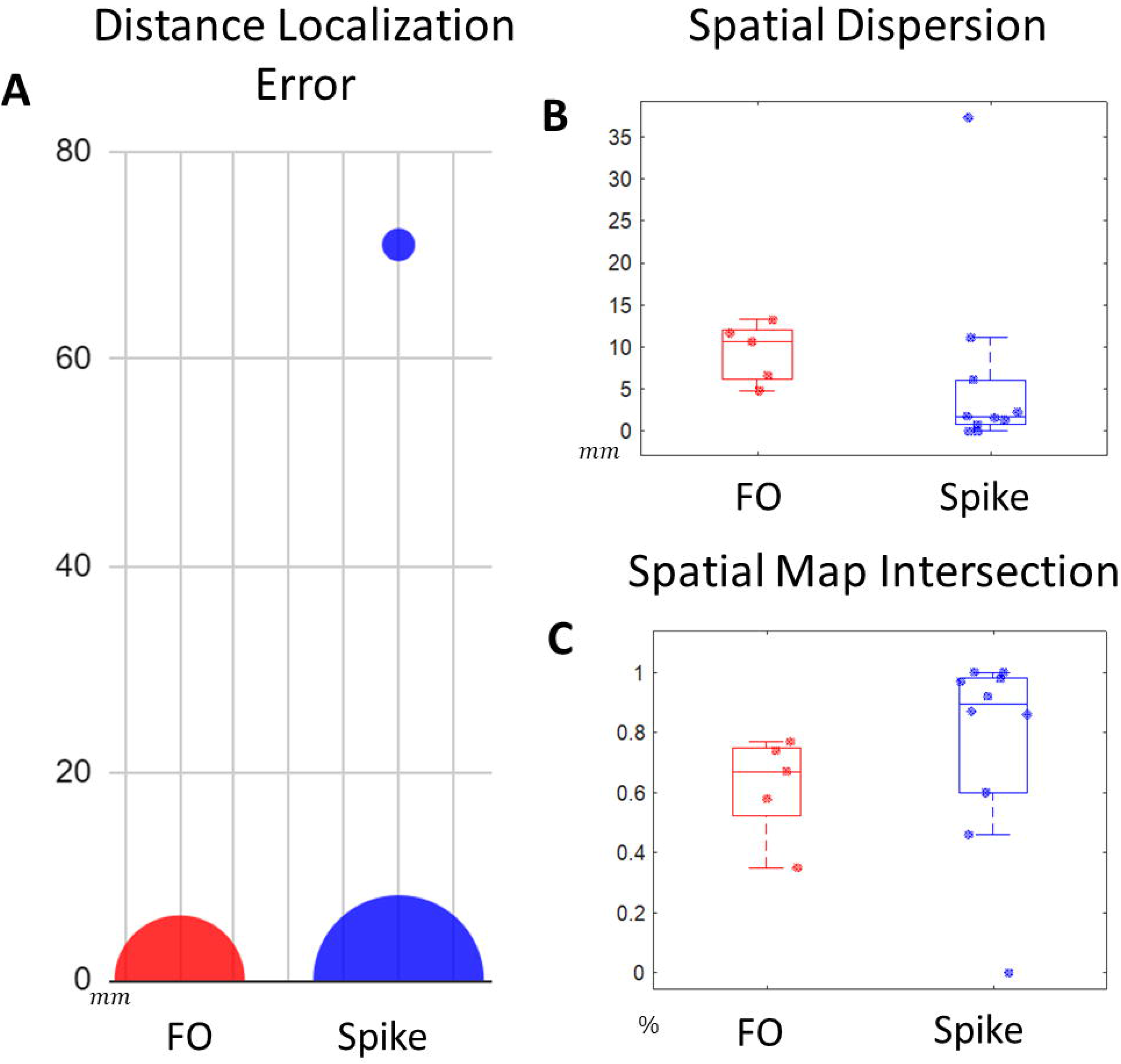
Validation metrics of the results of electrical source imaging of fast oscillations (FO) and spikes. Different measures were used to evaluate the success rate of the detectability and localization of FO compared to spikes (A) Distance Localization Error (DLE) (Dmin) as measured from the maximum of the localization map to the closest point of the cavity (Dmin=0 mm means within cavity; the larger the number, the further is the localization peak outside the cavity) (B) Spatial dispersion (SD) maps (50% threshold) measuring the amount of activity outside the cavity in relation to the distance from the closest cavity point (C) Spatial Map Intersection (SMI) between the activation maps (50% threshold) with the cavity such that 100% means that the localization map is completely localized inside the cavity.

### 3.3 Usefulness of the consensus map approach for FO source localization

The consensus map approach for FO sources resulted in less disperse maps compared to the one obtained by simply averaging of all FO maps (SD difference between cluster versus averaged map: 10.3±4.8mm versus 9.4±3.2mm; p=0.4). This difference in SD became significant when no thresholding was considered prior to the estimation of SD, resulting in a SD of 18.1±3.1mm for the averaged map compared to 16.3±3.2 mm for the consensus map (p=0.007). Our clustering approach was used instead of traditional averaging in order to have a more statistically robust localization. In the case of a small number of events as present for FO the clustering acts as a denoising mechanism as even one “bad” FO can misinform the whole average map. This demonstrated that the hierarchical clustering approach considered for the consensus map helped to remove the impact of more noisy FO events, resulting in a more reliable localization. This is illustrated in Figure. 6, which shows only the activations outside the cavity for averaged versus consensus FO source maps, when no threshold was applied.

**Figure 6.**
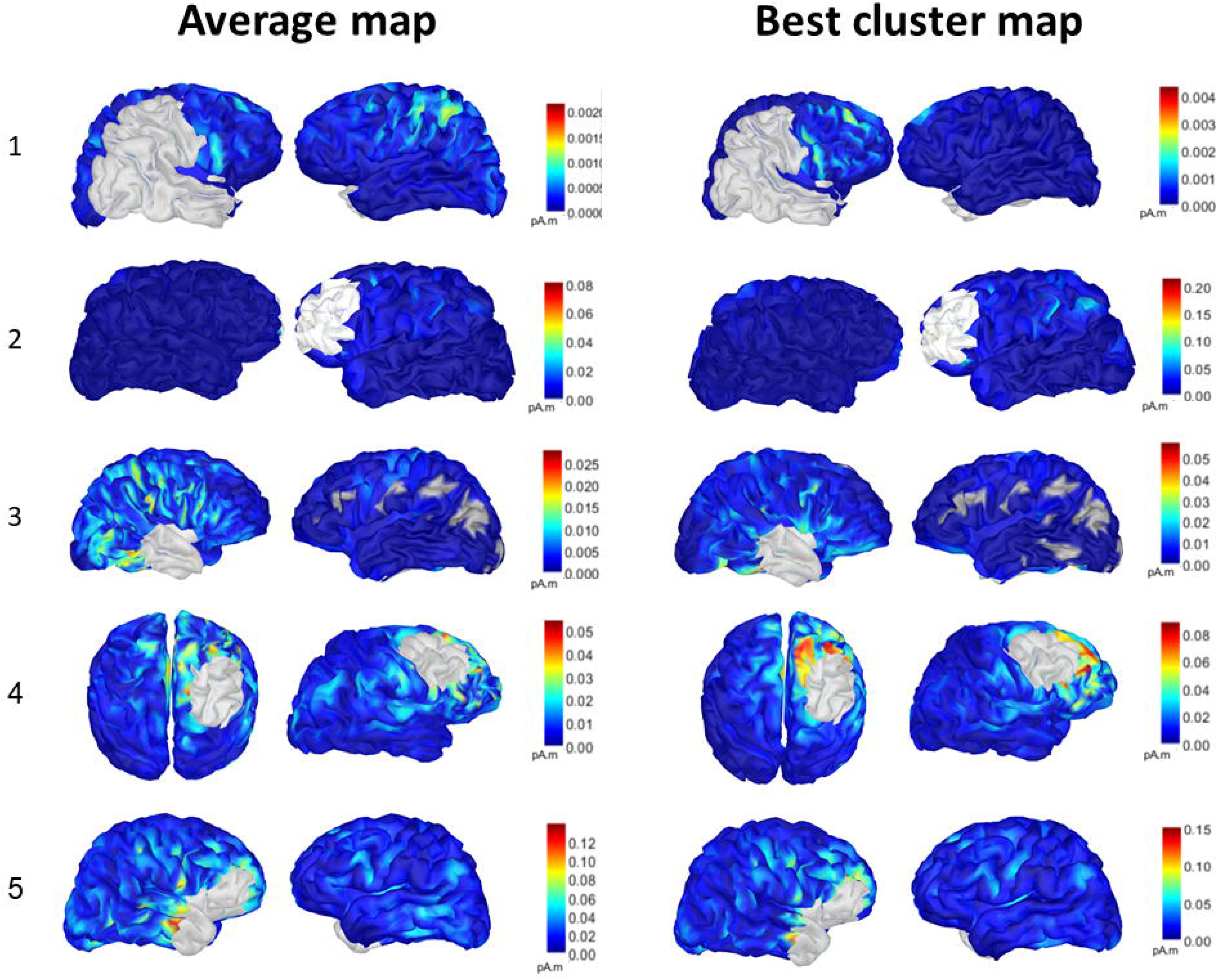
Comparison between electrical source imaging performed with the average vs. the unthresholded consensus map for fast oscillations (FO). FO were clustered using hierarchical clustering. The cluster exhibiting the most events was chosen as a consensus map. The cavity is marked in gray and the spurious activity outside the cavity is then depicted. Note that the maximum is inside the obscured cavity (as shown in Fig. 3) and only the spread is shown; thus the maximum is obscured by the cavity.

### 3.4 FO localization performance in HD-EEG vs the conventional 10-10 & 10-20 EEG arrays

We examined the localization accuracy of FO using the 10-10 system with 73 electrodes and the 10-20 system with 25 electrodes (Figures 3, 7 Supplementary Table 1). When considering the 10-10 system, among the FO detected using the HD-EEG array, we could identify FO in 4 out of 5 patients in the 10-10 electrodes. For the 10-20 system, we could identify FO in 3 out of 5 patients. Among the FO detected, all of them exhibited sufficient energy in the time-frequency representation and were therefore localizable using our proposed strategy. Results showed that using the full HD-EEG array we obtained a maximum source accurately localized in the cavity (Dmin = 0mm) for all cases (Figure 7), whereas with the 10-10 and 10-20 system, one and two FO localization exhibited a non-zero Dmin in each case (respectively). This means that the 10-10 system managed to correctly identify the EZ using FO in 60% (3 out of 5 patients) and the 10-20 system in 40% (2 out of 5 patients) of patients when detection and FO localization was considered. SD which measures the off-target spatial spread around the surgical cavity was shown to be lower with HD-EEG compared to that of the 10-20 system in all cases and to that of the 10-10 system in 2/4 of cases. SMI which represents the amount of spatial overlap with the surgical cavity, was higher with HD-EEG compared to the 10-10 system and 10-20 system in 2 of 4 cases and 2 of 3 cases respectively. SD and SMI values for each patient are provided in Supplementary Table 1.

**Figure 7.**
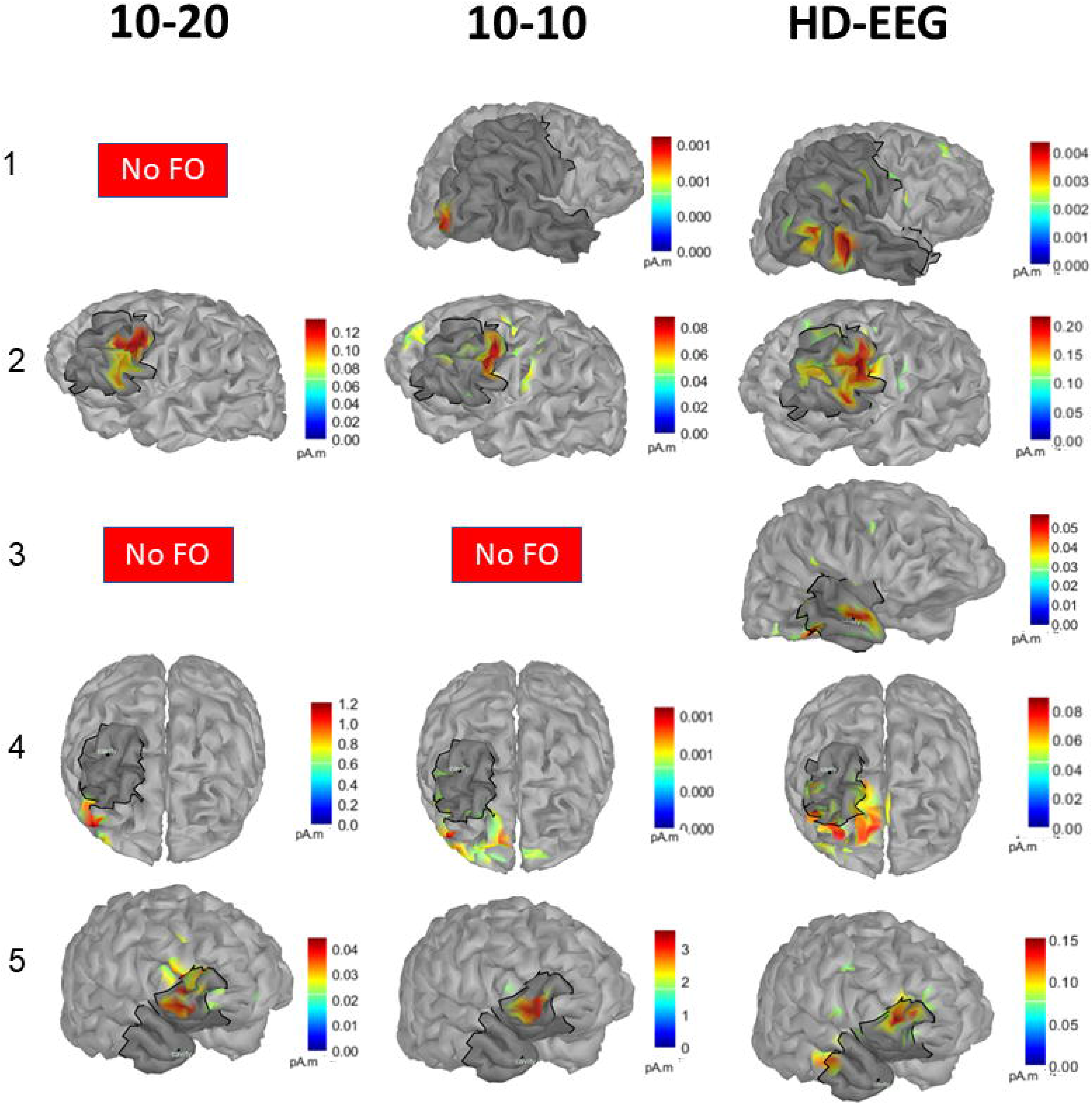
Results of electrical source imaging for fast oscillations (FO) for the 10-10 and the 10-20 electroencephalogram system. FO were source localized using the wavelet Maximum Entropy on the Mean method. The cavity is superimposed in light gray color. Final maps were either based on the best cluster or averaged in the case of < 8 FO events.

### 3.5 Generator depth

We found that all patients with localizable FO actually had a surface close generator, with 4 patients being classified as having a superficial generator and 1 patient being classified as having an intermediate generator (Table 2). In the 5 patients not showing sufficient numbers of localizable FO, the generator proved to be deep in 4 cases and intermediate in 1 (p=0.02). In the patient in whom the spike map localized outside the surgical cavity, the generator was actually localized in the cingulate gyrus, which is a deep region usually not detectable from scalp EEG.

## 4. DISCUSSION

This study presents a proof-of-concept that electrical source imaging of FO, if present, is able to identify the EZ in patients with drug-resistant focal epilepsy using HD-EEG. We demonstrated that (i) FO can be recorded in scalp high-density EEG during short recording sessions lasting 90 minutes, (ii) the presence or absence of sufficient numbers of FO is dependent on the depth of the epileptic generator, and (iii) electrical source imaging of FO using HD-EEG is able to correctly localize the EZ being superior to conventional EEG approaches using the 10-10 or 10-20 EEG systems.

### 4.1 Use of FO for identification of the EZ

FO were present in 50% of the investigated patients despite only short segments with light sleep used for detection of FO. Interestingly all patients exhibiting FO and spikes had a surface close generator, whereas the patients in whom we did not identify FO but only spikes had an either intermediate or deep generator. This finding is in keeping with a previous study investigating high-frequency oscillations in the scalp demonstrating that scalp FO are correlated with the depth of the epileptic generator (Cuello Oderiz et al., 2017).

We found that the maxima of the electrical source localization maps of FO correctly localized the EZ as approximated by the surgical cavity in all patients. The ability of FO to non-invasively localize the EZ was recently demonstrated using MEG (von Ellenrieder et al., 2016; Nissen et al., 2016; Velmurugan et al., 2019; Yin et al., 2019; Tamilia et al., 2020). Velmurugan and colleagues (2019) have recently shown the utility of high-frequency oscillations in correctly localizing the EZ in MEG in 52 patients thus demonstrating its potential as a scalp biomarker. Yet, while MEG has advantages such as better SNR and less complicated head modeling impacting spatial resolution when compared to EEG (Ilmoniemi and Sarvas, 2019), the lack of availability in most centers and its high maintenance cost makes it less ideal as a tool for everyday use in presurgical evaluation. In addition, it is worth noting that while EEG has been shown to detect higher numbers of FO (van Klink et al., 2019, Tamilia et al., 2020), spatial coverage is key due to the low SNR and a high-density array is preferable. Thus, this proof of principle study using HD-EEG with 256 electrodes together with the already existing evidence in MEG and EEG (Thomschewsky et al., 2019; van Klink et al., 2019, Tamilia et al., 2020) points to HD-EEG being a feasible candidate for FO localization of the EZ during pre-surgical evaluation.

### 4.2 Comparison of FO to spike sources

When comparing FO to spike source localization, we found that FO source localization and spike source localization were concordant with the latter resulting in localizations in all 10 study subjects. Note is made that one of the spike localizations was outside and far from the epileptic generator. The patient was identified as having a deep generator in the posterior cingulate gyrus determined by a focal MRI lesion and positron emission tomography (PET) hypometabolism, and the spikes were localized along the cortical surface corresponded hence to propagated activity. In this patient, FO were absent suggesting that source localization results in presence of FO are very likely pointing to the “true” onset generator (Cuello-Oderiz et al., 2017). Future research will assess the clinical validity of these results. In order to assess the feasibility of using scalp FO, and their eventual occurrence when superficial generators are involved, as a marker of the spike onset zone, a larger cohort and longer recording durations are needed. Our finding might hence be able to address a longstanding problem inherent to non-invasive source localization. However, careful evaluation of propagation of source localization along the peak of the spike vs. FO localization was out of the scope of this study and will be considered in future investigations. A recent study showed that electrical source imaging seems to be more accurate when performed at the time of spike onset instead of the peak of the spike (Plummer et al. 2019)

### 4.3 Different measures for source localization quality

In clinical practice most studies assessed the concordance of the source with the assumed EZ at the lobar (Duez et al., 2019; Rampp et al., 2019) or sub-lobar level (Heers et al., 2016). We evaluated our findings using different quantitative validation metrics, several of them being similar to the ones proposed in Pellegrino et al. (2018, 2020) and Chowdhury et al. (2018) in order to assess the quality of our results as objectively as possible regarding the source map maxima, its spatial extent, and the spatial intersection with the resection cavity. The originality of our proposed approach was also to consider an accurate delineation of the resection cavity as our reference for the evaluation of source localization results, whereas such a comparison was more qualitative (Abdallah et al., 2017) or semi-qualitative in other studies (Pellegrino et al (2018, 2020). The accuracy of FO localization was high with the maxima localized to the surgical cavity in all patients. Albeit being not significant, the extent of the source seemed to be more widespread in FO compared to spikes. However, whereas the estimation of the spatial extent of spike maps with cMEM has been carefully evaluated by our group (Chowdhury et al., 2016; Pellegrino et al., 2016; Pellegrino et al., 2020), the evaluation of the spatial extent of wMEM for FO would require further careful investigation. Moreover, it is difficult to disentangle whether the spatial extent of FO maps was true extent or only resulted from the localization of low SNR events. This might be potentially explained by the small number of FO given the short recording time of 90 minutes, as well as by the lack of consolidated sleep in the present study; longer recording durations preferably overnight are likely to be more favorable regarding FO quantity and SNR. Future research performing prolonged overnight sleep recordings is awaited for confirmation.

### 4.4 FO localization performance with different EEG montages

We confirmed evidence of previous work that a higher and denser electrode coverage is preferable to detect FO given their small generators (Kuhnke et al., 2018). Moreover, we demonstrated that this also impacts FO source localization. We showed that the correct localization of the EZ declines from 100% with HD-EEG to 60% with the 10-10 system and 40% with the 10-20 system. The use of HD-EEG seems therefore to be even more important for FO than spikes due to a lower SNR (Song et al., 2015). Further research with a larger cohort is needed in order to clinically validate our method for epilepsy patients undergoing surgery and see how our localization algorithm will perform.

### 4.5 Importance of consensus map approach for FO source maps

In this study we applied for the first time for spike and FO in HD-EEG the consensus map approach we recently proposed as a more robust approach than events averaging to provide reliable source localization, while taking into account the reproducibility of single discharges source maps (Chowdhury et al., 2018). In this study, this was our first attempt to consider consensus maps from wMEM results for FO localization. We therefore first performed a single event FO source localization, using only the TF box exhibiting the largest amount of energy along the FO, following the exact same methodology proposed in MEG by our group (von Ellenrieder et al., 2016; Chowdhury et al., 2018). We then compared every single FO source map using a hierarchical clustering in order to separate the data from events that were not in agreement with the majority of events and which would therefore add noise to the maps, as done in our previous work for spikes (Chowdhury et al., 2018). This consensus map approach seems to be particularly useful for FO, which in case of a discordant FO event tend to create a noisier map. Figure 4 shows a significant improvement of source localization of FO using this consensus map approach, when compared to standard averaging of all FO maps, resulting in a reduction of the SD of the source map outside the presumed EZ as approximated by the surgical cavity. Whether the fact of a lower accuracy of 0.87 in a recent combined EEG/MEG study (Tamilia et al., 2020) can be explained by the advantage that a consensus map could offer awaits further confirmation.

### 4.6 cMEM and wMEM for source localization of epileptic discharges

Most previous work in source localization of FO in MEG used the Beamformer technique, as it is assumed to be able to detect distributed and deep sources (Hu et al., 2017). Yet Beamformer is not a source localization method in its proper sense, but corresponds rather to a statistical dipole scanning approach, iteratively assessing how likely it would be to fit an equivalent current dipole at a specific position in a 3D grid covering the brain (Hillebrand et al., 2005). One important feature of the Beamforming technique is its inherent denoising properties (Cheyne et al., 2007; Hillebrand et al., 2005), as a spatial filtering approach, which is probably the main reason why several groups have considered this localization approach for FO (Belardinelli et al., 2012; van Klink et al., 2018). In this study, we chose the MEM framework, as one of the only distributed approaches that has been proposed to carefully recover the generators of epileptic discharges together with their spatial extent (Birot et al., 2011; Chowdhury et al., 2016). We also considered the time-frequency based extension of MEM (wMEM), providing us with the unique property of localization in the frequency domain (Lina et al., 2014), thus making it a good tool for the localization of oscillatory events, as previously demonstrated for FO in MEG and low-density EEG (von Ellenrieder et al., 2016; Tamilia et al., 2020), localization of oscillatory patterns at the seizure onset in EEG and MEG (Pellegrino et al., 2016; Pellegrino et al., 2018) and ongoing MEG resting state fluctuations (Aydin et al., 2020). For future investigations, it might be interesting to use a combined Beamformer-wMEM approach in order to increase the chance of FO detection with Beamformer applied as a denoiser in the source space followed by localization of the FO from denoised scalp data using wMEM.

### 4.7 SNR as a technical challenge

The SNR of EEGs can vary widely with different conditions as it is very susceptible to muscle artifact notably as well as interferences with outside noise (Islam et al., 2016). This latter challenge might be potentially overcome by the use of a low-noise amplifier (Fedele et al., 2015), which could make it possible to obtain low noise recordings in the future. We attribute the larger spread of FO maps compared to spike maps to the difference in the respective SNR which was shown to create a less focused localization for oscillatory activity with no spatial smoothing (Lina et al., 2014). The low SNR might also explain the fact that we detected mainly FO in the gamma frequency range (40-80 Hz) and not FO > 80 Hz. Prolonged clean recordings can be extremely valuable for FO identification, as data have the least artifacts in the high-frequency domain during sleep (Zijlmans et al., 2017). Yet in the case of prolonged recordings it might not be feasible to visually mark and validate each FO and a computational approach might be needed (Höller et al., 2018). Another interesting aspect that might influence the results is the type of electrode being used. Recent research demonstrated that tripolar EEG electrodes might yield some benefit with higher signal quality (Toole et al., 2019).

### 4.8 Limitations

First, we would like to acknowledge that our sample size is small. However, all 10 patients selected for this study were postsurgically seizure-free with an at least 2-year postsurgical follow-up, which allowed us to have the best estimate possible for approximation of the EZ in this proof-of-principle study. An alternative would have been to validate against the intracranially identified SOZ (Papadelis et al., 2016; Kuhnke et al., 2018; Dirodi et al., 2019). Given however, that up to ~50 % of patients in whom the SOZ has been surgically removed, do not become seizure-free after surgery (Krucoff et al., 2017), this seemed to us to not be the best option for the validation of our approach, even though a large resection diminishes the sensitivity to the real EZ (see patient 1 for an example). Second, we only had short recordings of approximately 1.5 h, during which patients achieved mostly only light sleep. Prolonged recordings of overnight sleep as planned in future research as well as use of denoising techniques as possible with Beamformer (Cheyne et al., 2007; Hillebrand et al., 2005) could result in higher patient numbers in whom FO can be identified and results regarding the extent of the source might be improved given higher event numbers and therefore better SNR. Third, we marked FO at the time of spikes as suggested by other authors (Nissen et al., 2016; Velmurugan et al., 2019), as FO at the time of spikes have a greater chance of a correct localization than FO occurring independent of spikes (Dirodi et al., 2019). To avoid a confound with a filtering effect of spikes (Bénar et al., 2010), we followed a two-step procedure to exclude false positives based on both visual signal inspection and presence of an isolated peak in the time-frequency representation in order to avoid misclassification of FO.

This study demonstrated the ability of 256-channel HD-EEG to correctly identify the EZ using source localization of FO. Presence or absence of FO was shown to be dependent on the presence of a surface close generator. This points to an added value of FO source localization, as presence of concordant spike and FO sources could confirm correct localization of the EZ, whereas lack of FO sources might point to the fact that the identified spike source could be the correlate of rather propagated activity and not the primary source, a problem inherent to non-invasive source imaging. We further confirmed the added value of HD-EEG for both FO detection and source localization, when comparing our findings to conventional approaches using the 10-10 and 10-20 EEG system. Further research is required to assess the clinical validity of these results. In order to assess the feasibility of using scalp FO in clinical practice, a larger cohort which includes successful and unsuccessful surgery outcomes is required to address the sensitivity and specificity of scalp FO in localization of the EZ and to compare their accuracy with that of spikes as traditional biomarker of epilepsy.

## ACKNOWLEDGMENTS

The authors wish to express their gratitude to Jean-Marc Lina, PhD, Professor at the École de Technologies Supérieures in Montreal for invaluable discussion of the wMEM framework used in this study for FO source localization. This work is supported by the Department of Medicine Innovation Fund 2016 of Queen’s University Internal Grant Competition to B.F. as well as a project grant of the Botterell-Howe-Powell Competition 2016 to B.F.. T.A.’s salary is supported by the Jenny Panitch Beckow Memorial Scholarship 2019-2020 of the Jewish Community Foundation of Montreal, start-up funding of the Montreal Neurological Institute to B.F. and a CIHR project grant PJT 159448 of C.G.. B.F.’s salary is supported by a salary award (“Chercheur-boursier clinicien Junior 2”) 2018 – 2021 of the Fonds de Recherche du Québec – Santé. AR acknowledges the support of the Italian Ministry of Health - Targeted Research Grant RF-2010-2319316.

