## Supplementary material for "Fast oscillations >40Hz localize the epileptogenic zone: an electrical source imaging study using high-density electroencephalography"

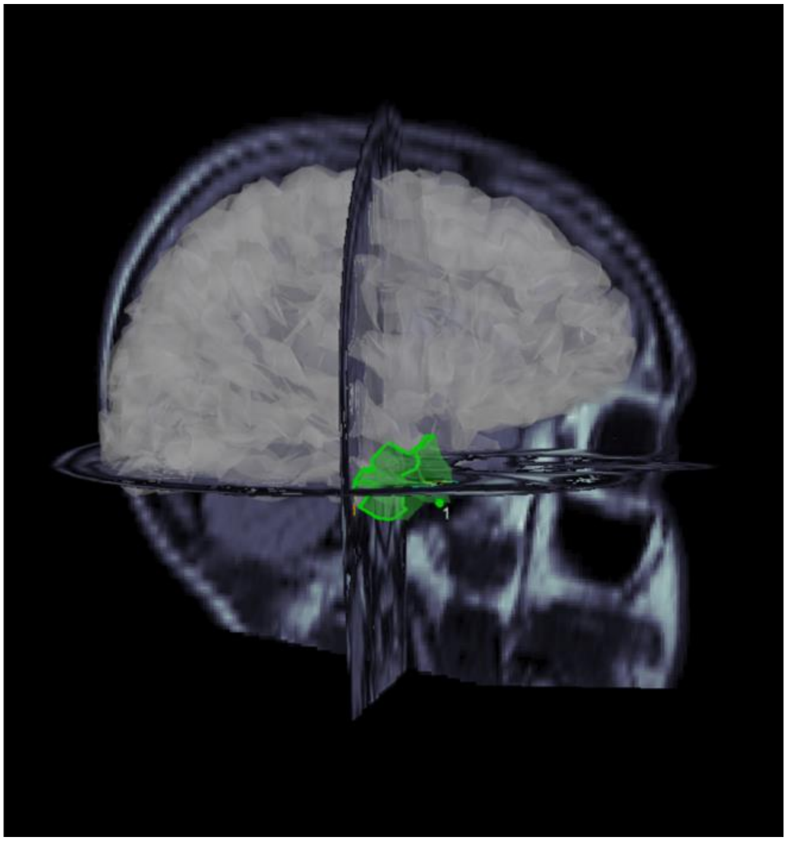

**Supplementary Figure 1.** Superimposing the scout of the cavity on the postsurgical MRI. The scouts were later used to determine the localization. The postsurgical MRI was co-registered into the space of the presurgical MRI using the MINC toolkit (https://bic-mni.github.io/). A 3D representation with the moveable surfaces was superimposed over the brain segmentation model. The brain was made semi-transparent in order to better visualize the superimposed post-surgical MRI. Once the resection cavity was identified, a scout was carefully fitted on the brain and grown until it filled the whole cavity. The scout was visually inspected and compared to the postsurgical MRI by a board certified neurologist.

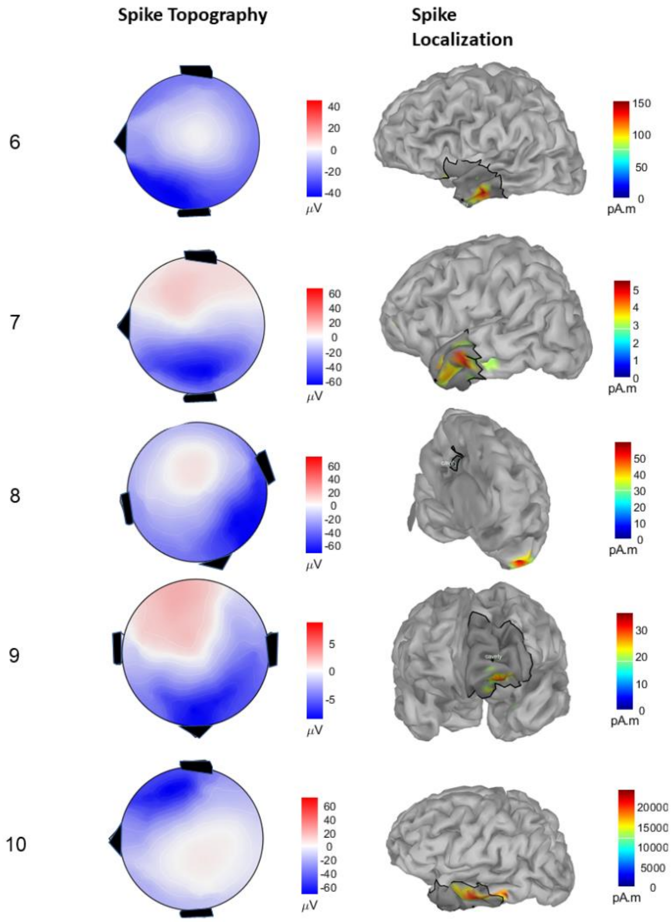


**Supplementary Figure 2.** Results of electrical source imaging of spikes for the 5 patients who did not have FO (see Fig. 3). Topography maps are provided on the left panel, source maps are provided on the right panel. The surgical cavity is superimposed on the brain model of the individual patients in dark grey color. Note that the topographical maps are oriented to the same direction as the source localization map results.

| # Patient | SD HD- EEG | SD 10-10 | SD 10-20 | SMI HD- EEG | SMI 10-10 | SMI 10-20 |
|---|---|---|---|---|---|---|
| **1** | 6.57 | 0 | - | 0.77 | 1 | - |
| **2** | 10.61 | 1.09 | 12.87 | 0.67 | 0.95 | 0.37 |
| **3** | 13.26 | - | - | 0.58 | - | - |
| **4** | 4.78 | 9.88 | 10.05 | 0.74 | 0.53 | 0.91 |
| **5** | 11.63 | 16.96 | 15.83 | 0.35 | 0.3 | 0.05 |

**Supplementary Table 1.** Fast oscillations (FO) localization metrices. The table shows Spatial Dispersion (SD) and Spatial Map Intersection (SMI) for the 256 electrodes high-density electroencephalogram (HD-EEG), 10-10 system, and 10-20 system. HD-EEG had a superior FO detection capability compared to the 10-10 system and the 10-20 system. Spatial dispersion (SD) which measures the off target spread was shown to be lower with HD-EEG compared to that of the 10-20 system in all cases and that of the 10-10 system in 50% of cases. Spatial Map Intersection (SMI) which represents the amount of overlap with the surgical cavity, was higher with HD-EEG in 67% of cases as assessed with the 10-20 system, and 50% of cases as assessed with the 10-10 system.
